# Efficacy of broflanilide (VECTRON™ T500), a new meta-diamide insecticide, for indoor residual spraying against pyrethroid-resistant malaria vectors

**DOI:** 10.1101/2020.11.06.367961

**Authors:** Corine Ngufor, Renaud Govoetchan, Augustin Fongnikin, Estelle Vigninou, Thomas Syme, Martin Akogbeto, Mark Rowland

**Author notes:** corresponding author Corine Ngufor.

## Abstract

The rotational use of insecticides with different modes of action for indoor residual spraying (IRS) is recommended for improving malaria vector control and managing insecticide resistance. A more diversified portfolio of IRS insecticides is required; insecticides with new chemistries which can provide improved and prolonged control of insecticide-resistant vector populations are urgently needed. Broflanilide is a newly discovered insecticide being considered for malaria vector control. We investigated the efficacy of a wettable powder (WP) formulation of broflanilide (VECTRON™ T500) for IRS on mud and cement wall substrates in WHO laboratory and experimental hut studies against pyrethroid-resistant malaria vectors in Benin, in comparison with pirimiphos-methyl CS (Actellic® 300CS). There was no evidence of cross-resistance to pyrethroids and broflanilide in CDC bottle bioassays. In laboratory cone bioassays, mortality of susceptible and pyrethroid-resistant *A. gambiae* s.l. with broflanilide WP treated substrates was >80% for 6-14 months. At application rates of 100mg/m^2^ and 150 mg/m^2^, mortality of wild pyrethroid-resistant *A. gambiae* s.l. entering treated experimental huts in Covè, Benin was 57%-66% with broflanilide WP and did not differ significantly from pirimiphos-methyl CS (57-66% vs. 56%, P>0.05). Mosquito mortality did not differ between the two application rates and local wall substrate-types tested (P>0.05). Throughout the 6-month hut trial, monthly wall cone bioassay mortality on broflanilide WP treated hut walls remained >80% for both susceptible and resistant strains of *A. gambiae* s.l.. Broflanilide shows potential to significantly improve the control of malaria transmitted by pyrethroid-resistant mosquito vectors and would thus be a crucial addition to the current portfolio of IRS insecticides.

**One Sentence Summary:** VECTRON™ T500, a new wettable powder formulation of broflanilide developed for indoor residual spraying, showed high and prolonged activity against wild pyrethroid-resistant malaria vectors, on local wall substrates, in laboratory bioassays and experimental household settings in Benin.

## Background

Indoor residual spraying (IRS) has historically been shown to be a powerful malaria control intervention (*1*). When applied correctly, IRS can quickly reduce malaria transmission by reducing adult mosquito vector density and longevity. It involves the application of a residual insecticide formulation to potential resting surfaces for malaria vectors such as internal walls, eaves and ceilings of houses, giving opportunity for vector mosquitoes to contact the insecticide and be killed in the process. IRS contributed substantially to the success of the malaria eradication campaign of the 1950s and 60s which resulted in the elimination of malaria from Europe and several countries in Asia and the Caribbean (*2*). The recent reductions in malaria morbidity and mortality observed in endemic countries in Africa and Asia over the last two decades has also been partly attributed to a significant increase in coverage with IRS (*3-5*).

The efficacy of IRS for malaria control is unfortunately threatened by widespread resistance in malaria vectors to the rather limited collection of insecticides approved for public health use (*6*). Pyrethroid resistance is now established across Africa and is increasing substantially in intensity the more they are used, making this previously ideal class of vector control insecticide almost unusable for IRS. Resistance to carbamates and organophosphates, which were for many years the only alternative IRS insecticide classes to pyrethroids (*5, 7*) is also increasing rapidly in malaria vector populations in Africa (*6, 8-11*). To mitigate the impact of insecticide resistance on malaria control, vector control programmes are encouraged to implement a rotational application of insecticides for IRS, alternating between insecticides with different modes of action (*12*). The use of rotations for insecticide resistance management relies on the concept that removing selection pressure for a given insecticide by switching between different modes of action, will result in resistance declining over time. An IRS rotation plan which will effectively reduce selection pressure for existing insecticide resistance genes and prevent the development of further resistance will, however, require a more diversified portfolio of IRS insecticides with more novel modes of action than what is currently available (*13*). This is driving the development of a new generation of IRS insecticide formulations containing new chemistries which can provide improved and prolonged control of insecticide-resistant malaria vector populations (*14-16*).

Broflanilide is a novel insecticide discovered by Mitsui Chemicals Agro, Inc (*17*) which has been formulated as a wettable powder for IRS. It has a unique chemical structure characterized as a *meta*-diamide which acts as a non-competitive antagonist of the γ -aminobutyric acid (GABA) receptor of chloride channels of the insect inhibitory nervous system (*18*). Broflanilide was classified by the Insecticide Resistance Action Committee (IRAC) as a GABA-gated chloride channel allosteric modulator (IRAC Group 30), causing hyperexcitation and convulsion in insects (*19*). Its mode of action is distinct from that of conventional non-competitive antagonists (NCAs) of the GABA-gated chloride channel, such as picrotoxinin, dieldrin, fipronil, lindane and α-endosulfan (*20*). There is no known cross-resistance between broflanilide and current public health insecticides. The active metabolite exhibits high selectivity for the insect RDL GABA receptor compared to the mammalian receptors (*18*) but exhibits no cross-resistance to dieldrin. Broflanilide has demonstrated excellent insecticidal activity against many insect species including Lepidopteran and Coleopteran pests and Thysanopteran pests (*17*) and has also shown low acute toxicity against non-target aquatic organisms (*21*) demonstrating high potential for public health and agricultural use.

In this study, we investigated the potential of VECTRON™ T500, a wettable powder (WP) formulation of broflanilide (broflanilide WP), for indoor residual spraying against mosquito vectors of malaria. The insecticide was assessed for its efficacy and residual activity on local IRS wall substrates in a series of WHO phase I laboratory bioassays with susceptible *Anopheles gambiae* s.s. and resistant strains of *A. gambiae* s.l. and in WHO phase II experimental huts studies against wild free-flying pyrethroid-resistant of *A. gambiae* s.l. in Southern Benin, West Africa.

## Results

### WHO Phase I laboratory bioassays

Following WHO guidelines (*22*), laboratory bioassays were performed to investigate possible cross-resistance to broflanilide and pyrethroid-resistance mechanisms in CDC bottle bioassays using technical grade insecticide and to identify an effective dose of broflanilide WP (VECTRON™ T500), for IRS using WHO cone bioassays. Cone bioassays were also performed to investigate the residual efficacy of broflanilide WP on cement and mud block substrates. The bioassays were conducted using laboratory maintained mosquitoes of the susceptible *A. gambiae* s.s. Kisumu strain and the pyrethroid-resistant *A. gambiae* s.l. Covè strain which has shown over 200 fold resistance to pyrethroids mediated by a high frequency of the knockdown resistance L1014F allele (>90%) and overexpression of the cytochrome P450 CYP6P3, associated with pyrethroid detoxification (*23*).

### No evidence of cross-resistant to broflanilide and pyrethroids in CDC bottle bioassays

Mosquito mortality rates following exposure in CDC bottles coated with alpha-cypermethrin 12.5µg were 100% with the insecticide susceptible *A. gambiae* s.s. Kisumu strain and 45% with the pyrethroid-resistant *A. gambiae* s.l. Covè strain thus confirming the high levels of pyrethroid resistance in the Covè strain (*23*). The mortality results of both strains exposed to broflanilide in the CDC bottles treated with a range of doses between 5µg and 200µg per bottle are presented in Figure 1. Using log dosage-probit mortality analysis, the lethal concentration (LC) required to kill 50% (LC50) and 95% (LC95) of exposed mosquitoes were 8.5µg and 70µg respectively with the susceptible Kisumu strain and 18.1µg and 73.6µg with the pyrethroid-resistant Covè strain (Table 1). A small resistance ratio of 2.1 (95% limits: 1.7-2.7) for the Covè strain to the Kisumu strain at the LC50 was thus detected suggesting the absence of cross-resistance to broflanilide and pyrethroids.

**Table 1:** Lethal dose of broflanilide on pyrethroid-resistant *A. gambiae* s.l. Covè exposed in CDC bottle bioassays

| Mosquito strain | Slope (SE) | LD50*<br>(95% CI) | LD95*<br>(95% CI) | Resistance Ratio<br>(95% CI) |
| --- | --- | --- | --- | --- |
| Susceptible *A. gambiae* s.s. Kisumu | 2.7 (0.14) | 8.5 (2.2-14.1) | 70 (38.5-141.1) | - |
| Pyrethroid-resistant *A. gambiae* s.l. Covè | 1.8 (0.20) | 18.1 (10.7-25.4) | 73.6 (46.9-214.9) | 2.1 [1.7-2.7] |

\*lethal doses are expressed in µg/ml

**Figure 1.**
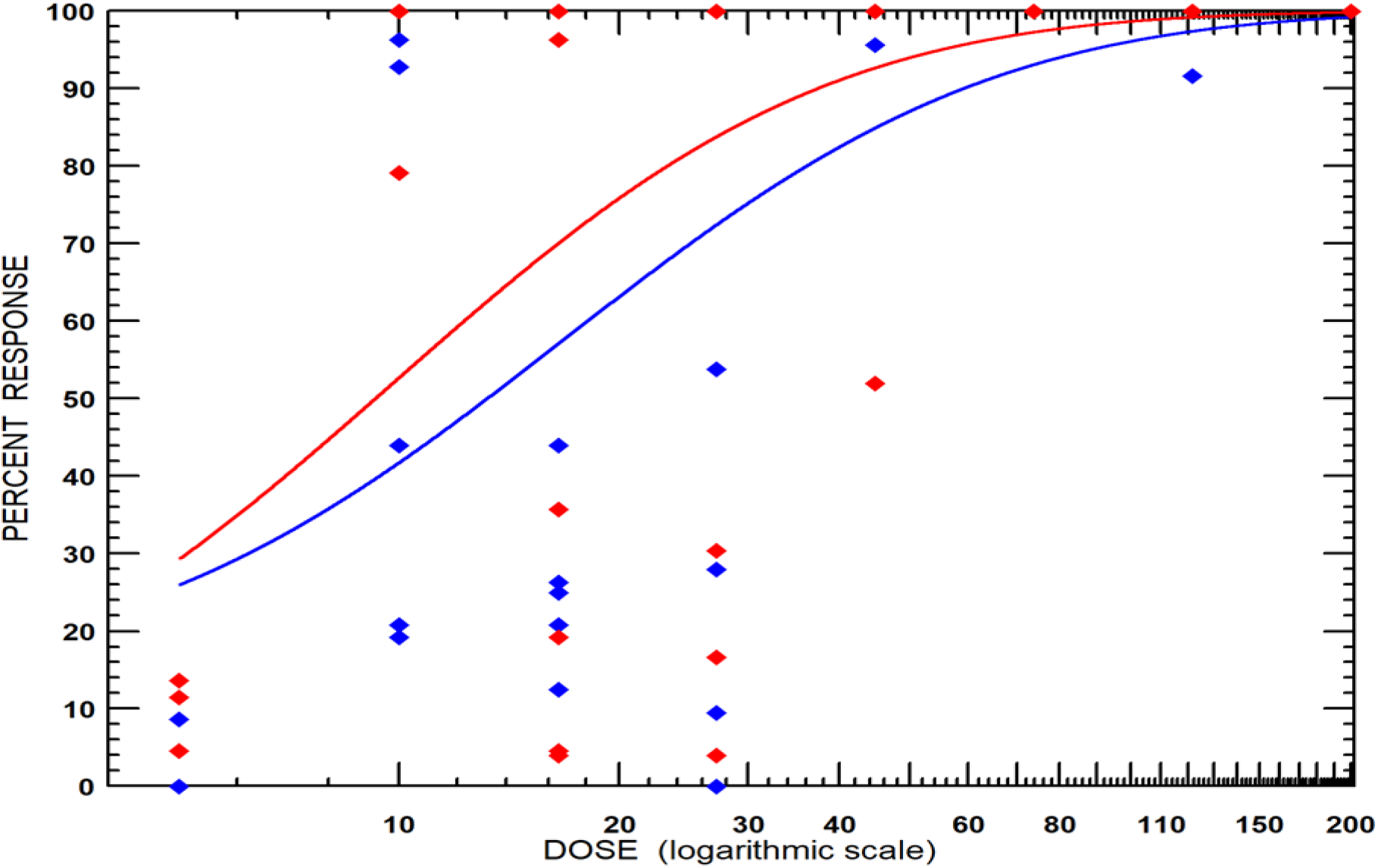
Mortality of susceptible *A. gambiae* Kisumu (red) and pyrethroid-resistant *A. gambiae* s.l. Covè (blue) mosquitoes in CDC bottle bioassays treated with a technical grade of broflanilide insecticide. Mosquitoes (150/dose) were exposed for 1-hour in cohorts of 25 per bottle and mortality recorded after 72h based on preliminary findings of delayed mortality effect with broflanilide. The red line represents the response of the susceptible Kisumu strain while the blue line represents the response of the pyrethroid-resistant Covè strain.

**Figure 1:**
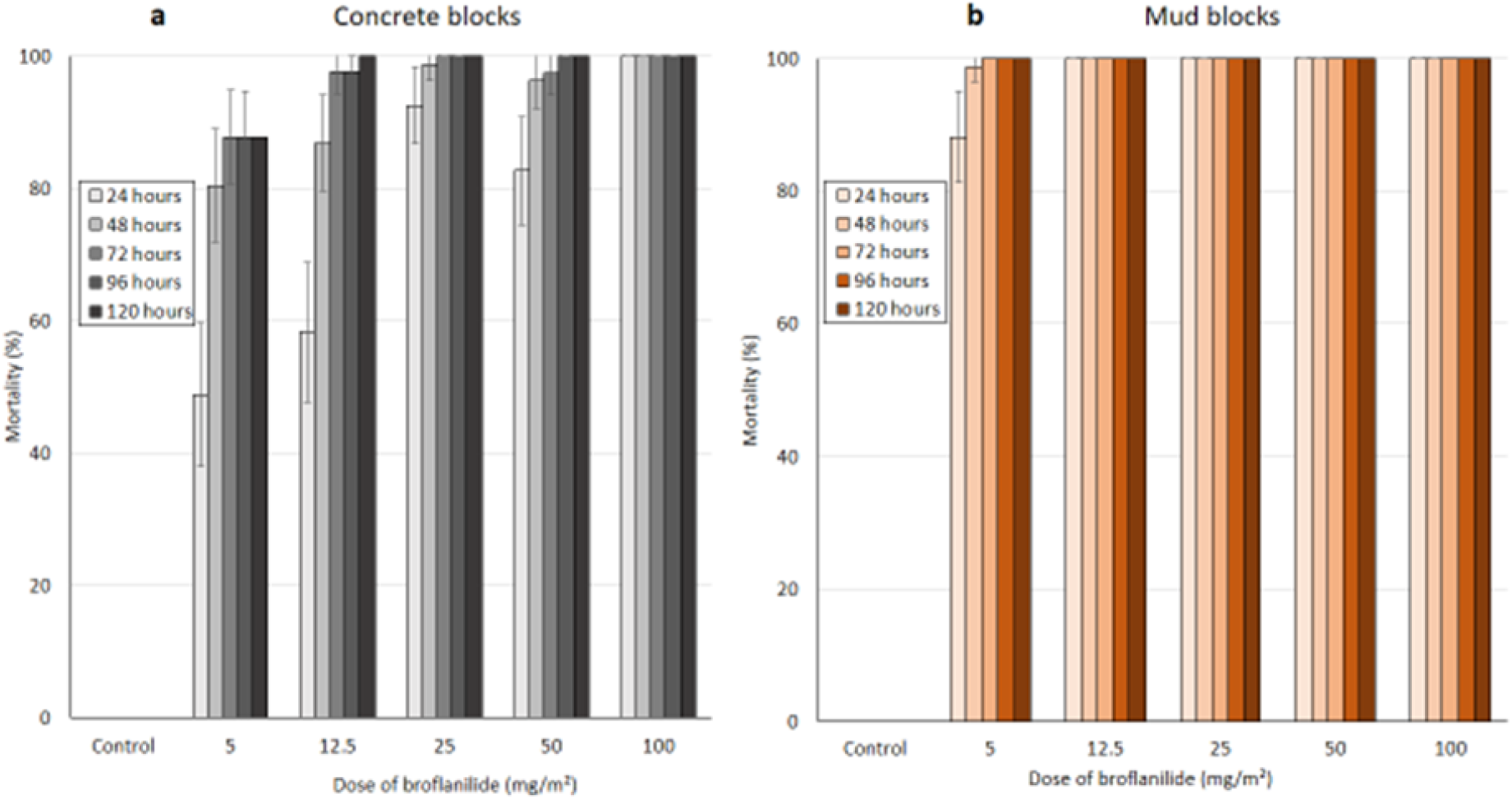
Mortality of insecticide-susceptible *A. gambiae* s.s. Kisumu strain mosquitoes on broflanilide WP treated cement (a) and mud (b) blocks substrates. Mosquitoes were exposed 1-week post-treatment for 30-minutes in WHO cone bioassays and mortality recorded every 24hrs for up to 120hrs.

### Dose-response cone bioassay studies

According to WHO guidelines (*22*), the target dose of a new insecticide for IRS should be investigated from doses 2-4 times the minimum dose that will cause 100% mortality in a fully susceptible mosquito vector population. To identify a suitable target dose of broflanilide WP for IRS, WHO cone bioassays were conducted on cement and mud block substrates treated with a range of concentrations of broflanilide WP between 5 mg/m^2^ and 100 mg/m^2^ to detect the minimum dose that will cause 100% mortality. The cone bioassays were performed 1-week post block treatment using the insecticide-susceptible *A. gambiae* s.s. Kisumu strain. The mortality rates observed in the cone bioassays are presented in Figure 1. No knockdown was recorded with broflanilide WP at any of the doses and with any of the substrates tested. Mortality reached 100% within 24 hours at a dose of 100 mg/m^2^ on cement block substrates (Fig1a) and a dose of 12.5 mg/m^2^ on mud block substrates (Fig 1b). Broflanilide WP performed better on mud than on cement. Data with cement blocks showed a delayed mortality effect with broflanilide WP doses below 100 mg/m^2^ with mortality increasing gradually from 24 hours and reaching a peak at 72 hours. At the doses tested, there was no measurable increase in mortality when holding time was extended beyond 72hours. This demonstrated a delayed mortality effect with broflanilide WP, as a result, for subsequent studies with the insecticide, mosquito mortality was recorded only up to 72hours post-exposure.

A dose of 200 mg/m^2^, which was two times the dose that induced 100% mosquito mortality within 24hours on both substrates, was identified as a suitable dose for further laboratory studies on the residual efficacy of broflanilide WP. Since the insecticide performed better on mud block substrates inducing optimal mortality at even much lower doses, residual efficacy on mud block substrates was also assessed at 100 mg/m^2^.

### Broflanilide WP shows prolonged residual efficacy on block substrates in laboratory cone bioassays

The residual efficacy of broflanilide WP was investigated at application rates of 200 mg/m^2^ on cement block substrates and 100mg/m^2^ and 200 mg/m^2^ on mud block substrates. Blocks of each substrate-type were treated at each selected dose and tested in WHO cone bioassays at 1-week post-treatment and monthly intervals subsequently using the insecticide-susceptible *A. gambiae* s.s. Kisumu and the pyrethroid-resistant *A. gambiae* s.l. from Covè.

Monthly cone bioassay mortality (72h) on cement blocks treated at 200mg/m^2^ was >80% with the susceptible *A. gambiae* s.s. Kisumu strain for 6 months after which it ranged between 57% and 100% up to month 18 post-treatment (Fig 2). With the pyrethroid-resistant *A. gambiae* s.l. Covè strain, mortality on cement blocks (200mg/m^2^) remained >80% for up to 14 months post-treatment (Fig 2). Monthly cone bioassay mortality of both strains at both doses tested on mud blocks (100 mg/m^2^ and 200 mg/m^2^) remained >80% for 16 months (Fig 3).

**Figure 2:**
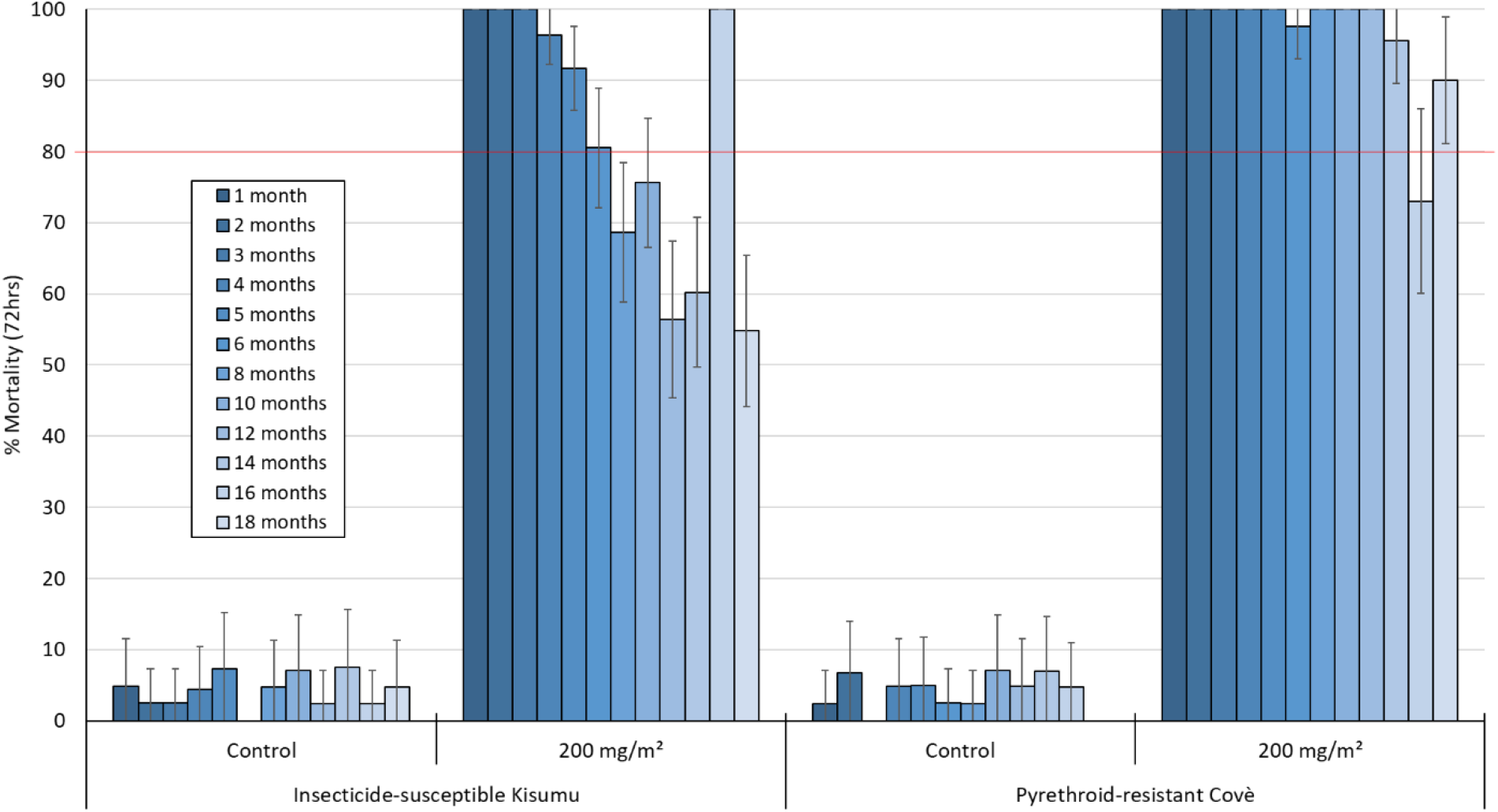
Monthly cone bioassays mortality of insecticide-susceptible *A. gambiae* s.s. Kisumu and pyrethroid-resistant *A. gambiae* s.l. Covè strain mosquitoes on broflanilide WP-treated cement block substrates in the laboratory (WHO Phase 1). At each time point, forty 2-5 days old female mosquitoes were exposed for 30-minutes in WHO cone bioassays and mortality recorded after 72h.

**Figure 3:**
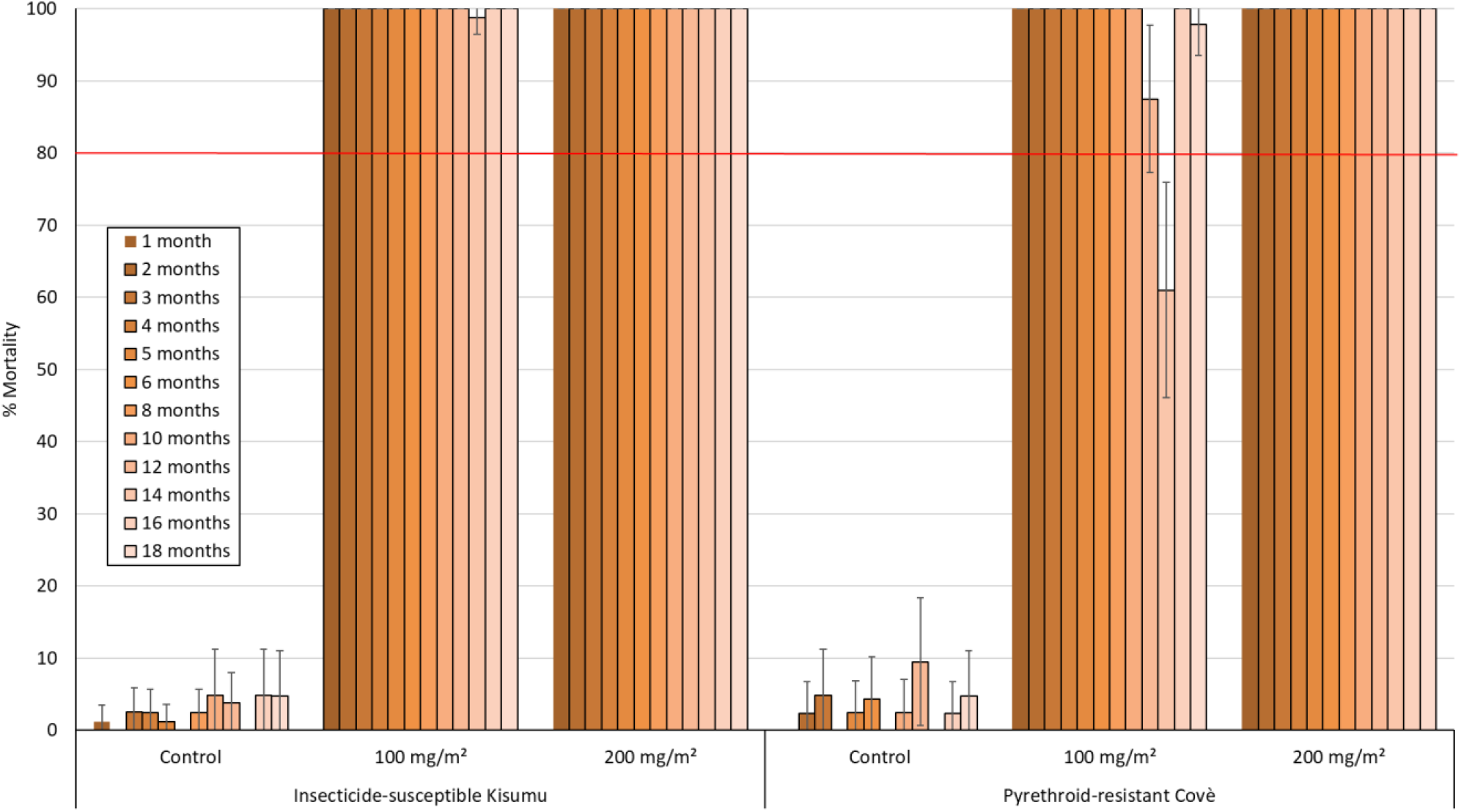
Monthly cone bioassays mortality of insecticide-susceptible *A. gambiae* s.s. Kisumu and pyrethroid-resistant *A. gambiae* s.l. Covè strain mosquitoes on broflanilide-treated mud block substrates. At each monthly time point, forty 2-5 days old female mosquitoes were exposed for 30-minutes in WHO cone bioassays and mortality recorded after 72h.

### WHO Phase II experimental hut evaluation of broflanilide WP in Covè, Benin

To investigate the efficacy of the broflanilide WP formulation for IRS against wild free-flying pyrethroid-resistant malaria vectors, we performed an experimental hut trial at the CREC/LSHTM experimental hut station in Covè, Southern Benin ([7°14′N2°18′E](https://geohack.toolforge.org/geohack.php?pagename=Cove&params=7_14_N_2_18_E_type:city(43554)_region:BJ)). The local vector population in Covè is resistant to pyrethroids and DDT. Molecular analysis has revealed a kdr (*L1014F*) allele frequency of 89% and microarray studies have also found overexpression of CYP6P3, a P450 that is as an efficient metabolizer of pyrethroids (*23*).

Experimental huts simulate household conditions and are thus used to assess the capacity of indoor vector control interventions to prevent mosquito entry, induce early exiting of vector mosquitoes, prevent mosquito feeding and induce mosquito mortality under carefully controlled household conditions (*22, 24*). Broflanilide WP was evaluated against wild pyrethroid-resistant *An gambiae* sl in Covè in experimental huts treated at application rates of 100 mg/m^2^ and 150 mg/m^2^. To investigate efficacy on commonly used wall substrates in Benin, for each application rate, the inner walls and ceiling of the experimental huts were plastered with either cement or mud. Broflanilide WP was compared to primiphos-methyl CS (Actellic® 300CS) applied at 1000mg/m^2^ on cement walls as a positive control. At the time of this study, the toxicity and potential risk of broflanilide WP for IRS was yet to be fully assessed, so it was not acceptable to use human volunteer sleepers as hosts to attract wild vector mosquitoes into the experimental huts until a human risk-assessment was performed and the product was found safe for IRS at the potential application rates. Preliminary studies revealed that the local wild *A. gambiae* s.l. at the Covè experimental hut station were also attracted to and blood-fed on cows, although less than to humans. Hence, cows were used as replacement hosts to attract mosquitoes into the experimental huts during the trial.

### Broflanilide WP induces high mosquito exiting rates

A total of 745 female *A. gambiae* s.l. and 771 female *A. ziemanni* were collected in the experimental huts over the 6-month trial (Tables 2 and 3). Molecular species analysis (SINE PCR) performed using the protocol proposed by Santolamazza *et al* (*25*) on DNA extracted from a random selection of 100 live and dead *A. gambiae* s.l. collected in the experimental huts during the 6-month trial, revealed that the vector was composed of 89% *A. coluzzii* and 11% *An gambiae* s.s.

**Table 2.**
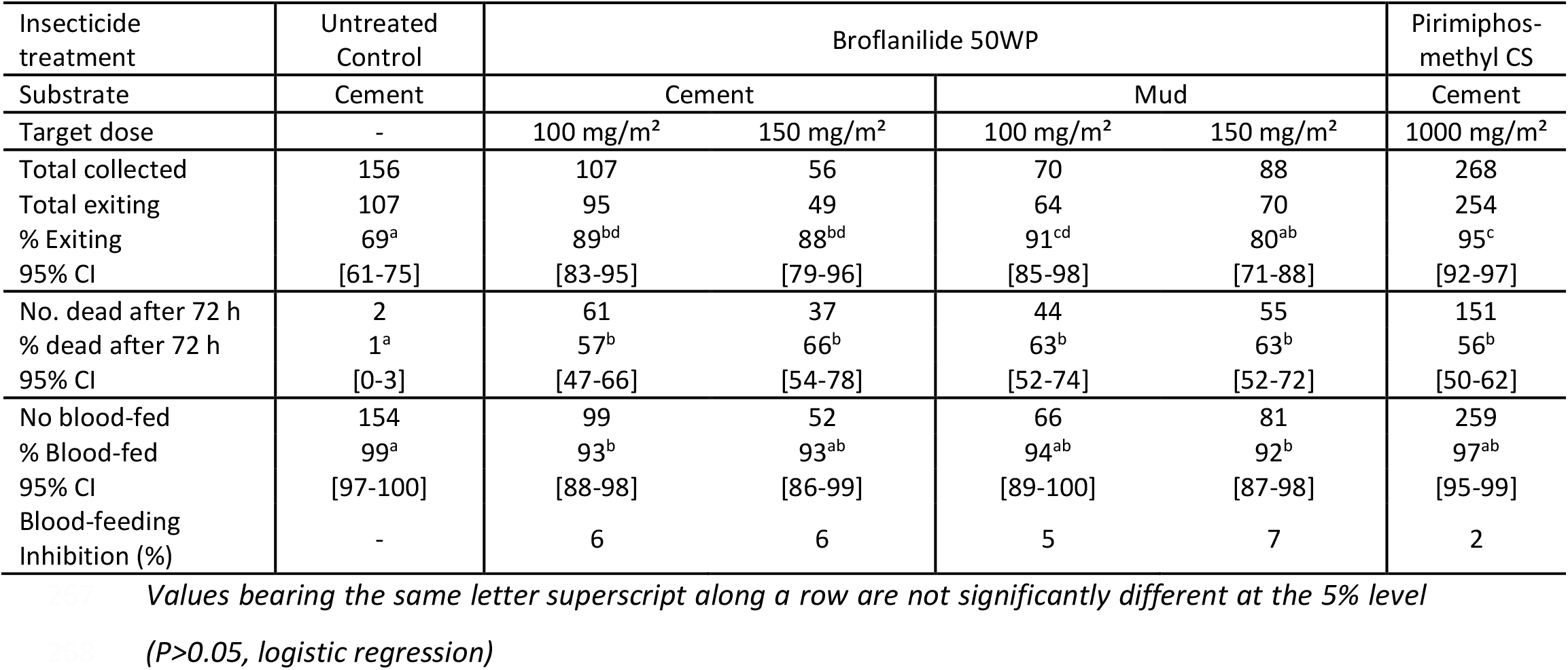
Results with wild, free-flying pyrethroid-resistant *A. gambiae* s.l. entering experimental huts in Covè, Benin

**Table 3.**
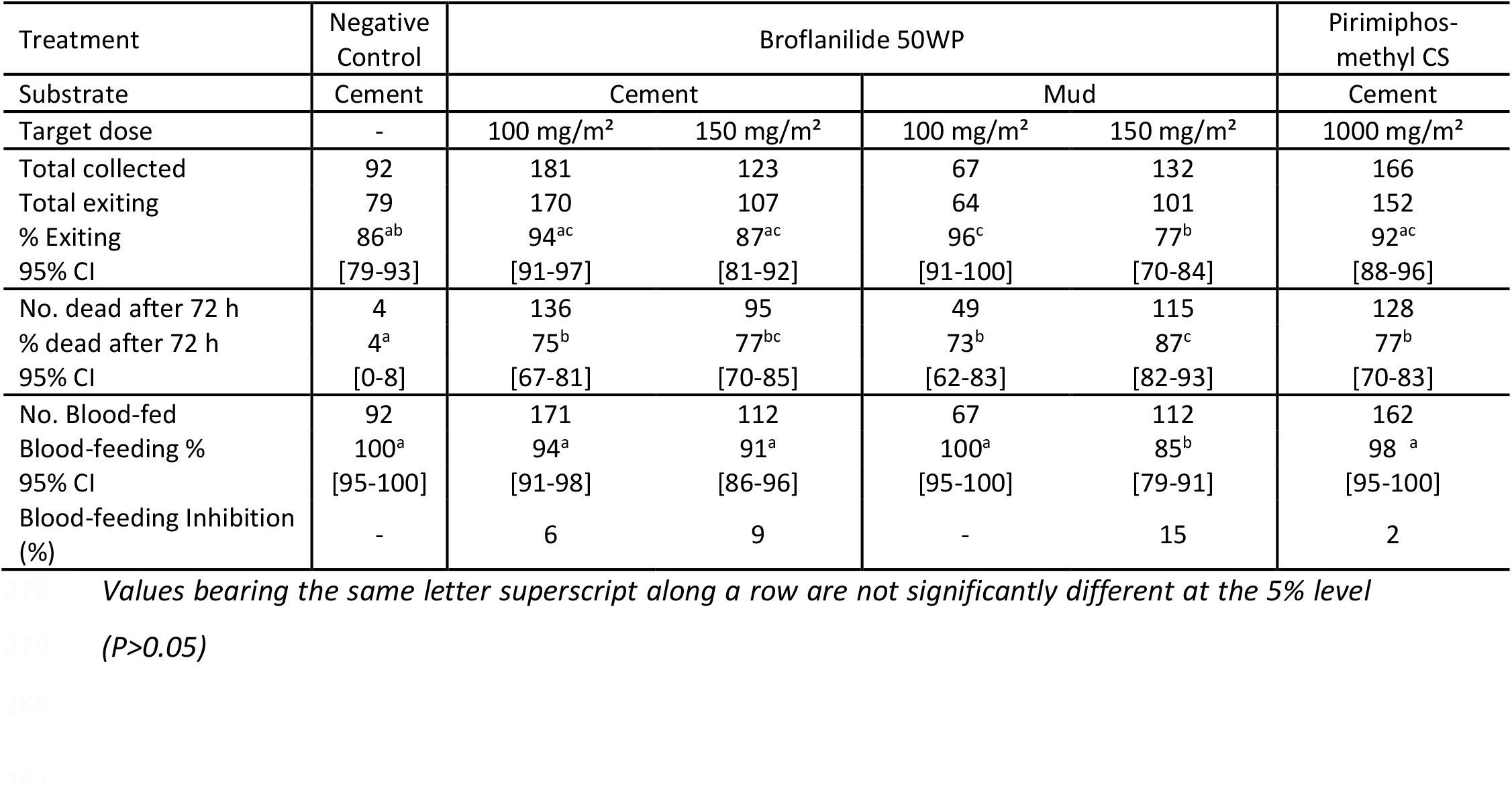
Results with wild, free-flying pyrethroid-resistant *A. ziemanni* entering experimental huts in Covè, Benin

Exiting rates of *A. gambiae* s.l. were generally higher in broflanilide WP-treated huts (88%-92%) compared to the control hut (69%), but only slightly lower than in pirimiphos-methyl CS-treated huts (95%). Exiting rates with broflanilide WP did not also differ substantially between the two application rates tested and between the two substrates assessed (Table 2).

A similar trend in mosquito exiting was observed with the *A. ziemanni* collected in the experimental huts except that exiting rates with broflanilide WP-treated huts did not differ significantly from that in the negative control hut (P<0.05) (Table 2).

### Broflanilide WP induces similar mortality of wild pyrethroid-resistant A. gambiae s.l compared. to pirimiphos-methyl CS

Mortality of free-flying wild, pyrethroid-resistant *A. gambiae* s.l. entering the control untreated hut was 1% while mortality with broflanilide WP treated huts ranged between 57% and 66% (Table 2). The mortality rates observed with broflanilide WP were similar to mortality observed with pirimiphos-methyl CS (57% - 66% with broflanilide WP vs. 56% with pirimiphos-methyl CS; P>0.05). For each substrate type, mortality with broflanilide WP did not differ significantly between the doses tested; cement: 57% with 100 mg/m^2^ vs 66% with 150 mg/m^2^ (P = 0.439) and mud: 63 % with both application rates (P = 0.922). Broflanilide WP generally performed the same on mud and cement substrates at both application rates; at 100 mg/m^2^: 57 % on cement vs. 63% on mud (P = 0.938) and at 150 mg/m^2^: 66% with cement vs. 63% with mud (P = 0.664). In addition, for each application rate, the difference in mortality between substrates was not significant (P>0.05). As in the phase 1 cone bioassays, the free-flying mosquito experimental hut data also showed a delayed mortality effect with broflanilide WP on *A. gambiae* s.l. (Figure 5). Mortality increased steadily from 35-41% at 24 hours to 57-63% at 72 hours with the 100 mg/m^2^ application rate and from 48% - 52% at 24h hours to 63% - 66% at 72 hours with the 150 mg/m^2^ application rate (P<0.05).

**Figure 5:**
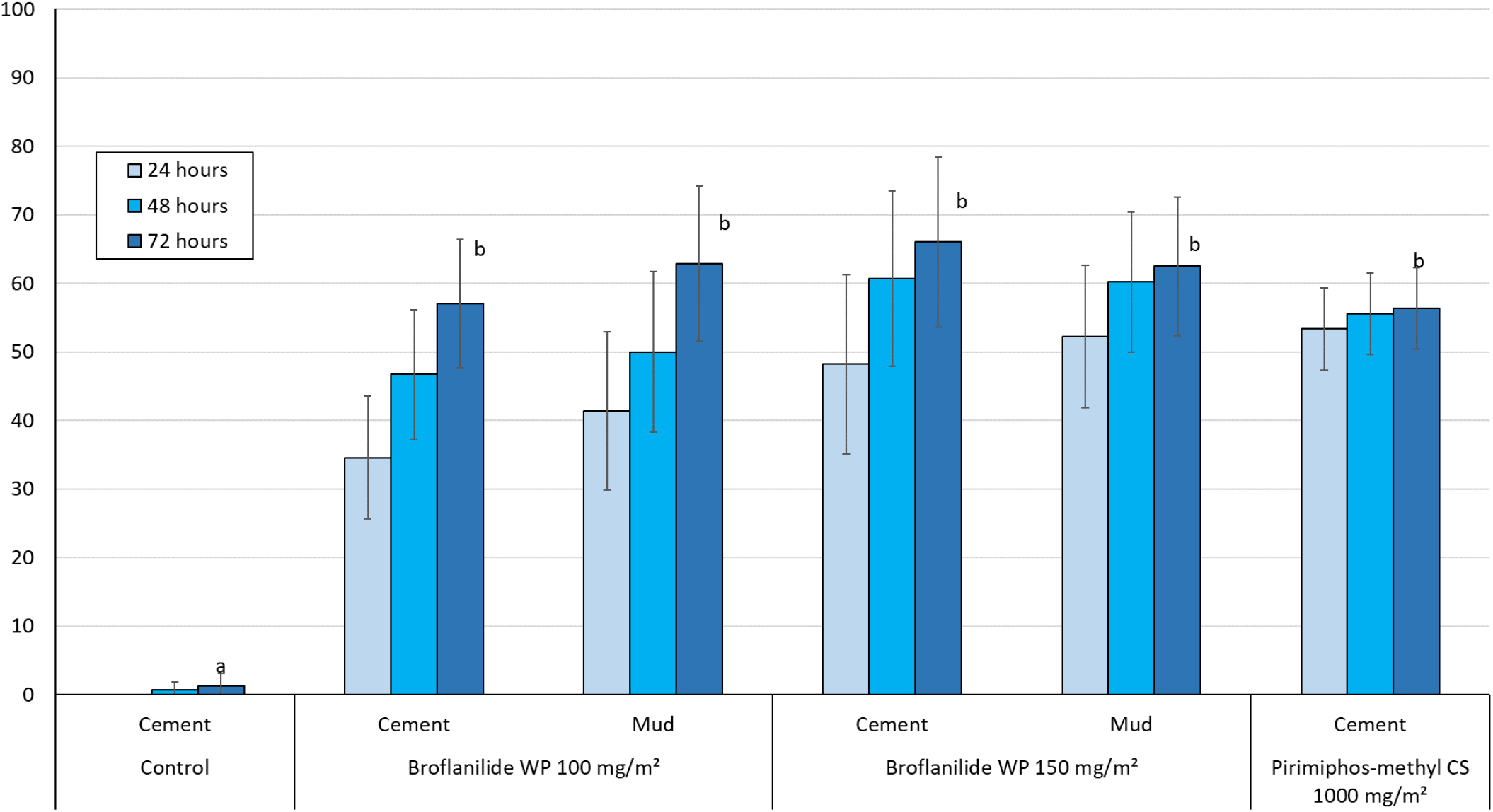
Overall mortality of wild free-flying pyrethroid-resistant *A. gambiae* s.l. at 24, 48 and 72 hours after collection from experimental huts in Covè, Benin. *Bars bearing the same letter label are not significantly different at the 5% level (logistic regression). Error bars represent 95% confidence intervals. Broflanilide WP induced a delayed effect on wild vector mosquitoes in experimental huts*.

**Figure 5.**
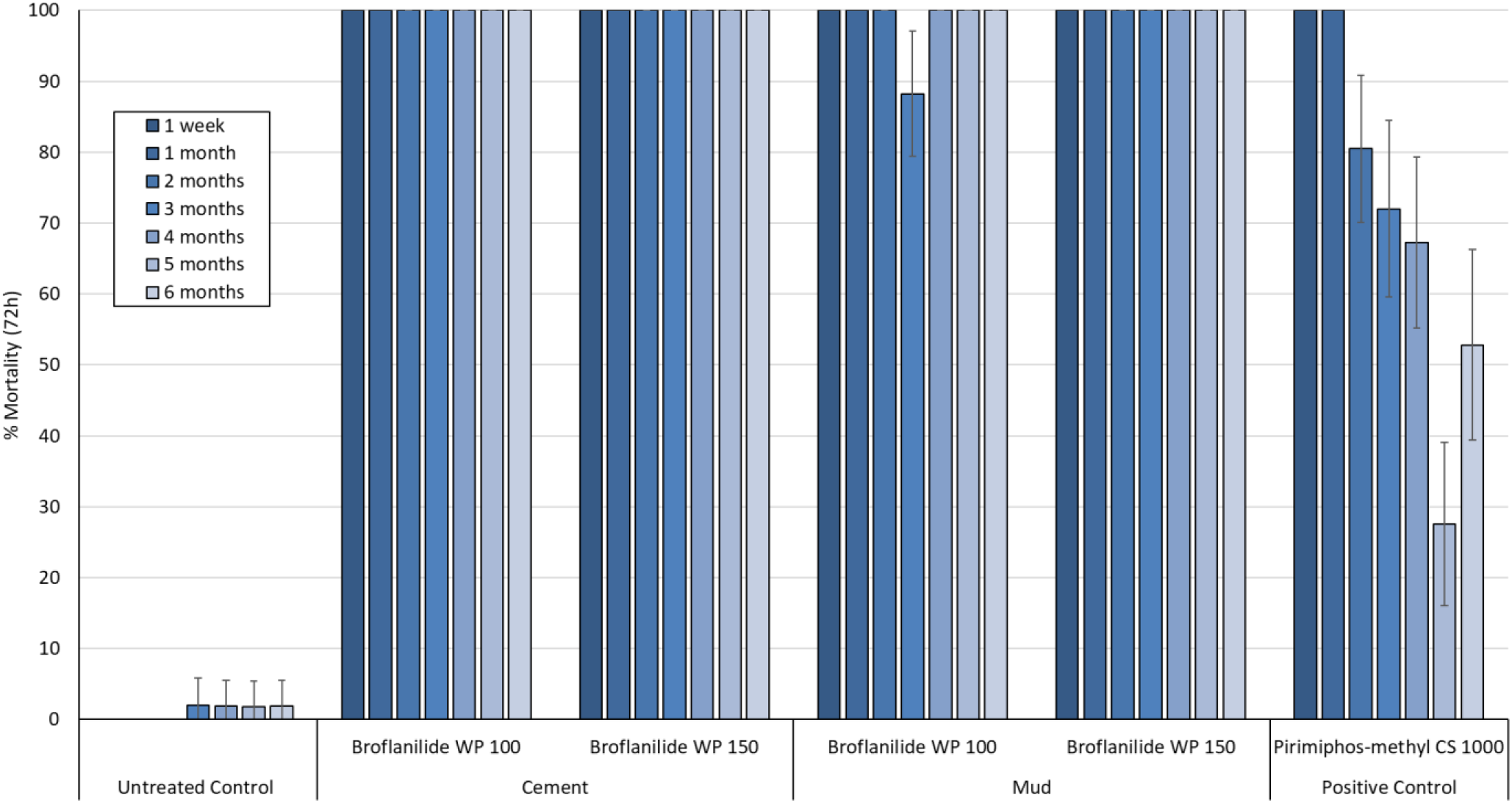
Cone bioassays mortality (72h) with susceptible *A. gambiae* s.s. Kisumu on broflanilide WP treated experimental hut walls. At each monthly interval, ∼50 2-5 days old female mosquitoes were exposed on treated of each hut in cohorts of 10 per cone. Error bars represent 95% confidence intervals.

**Figure 6.**
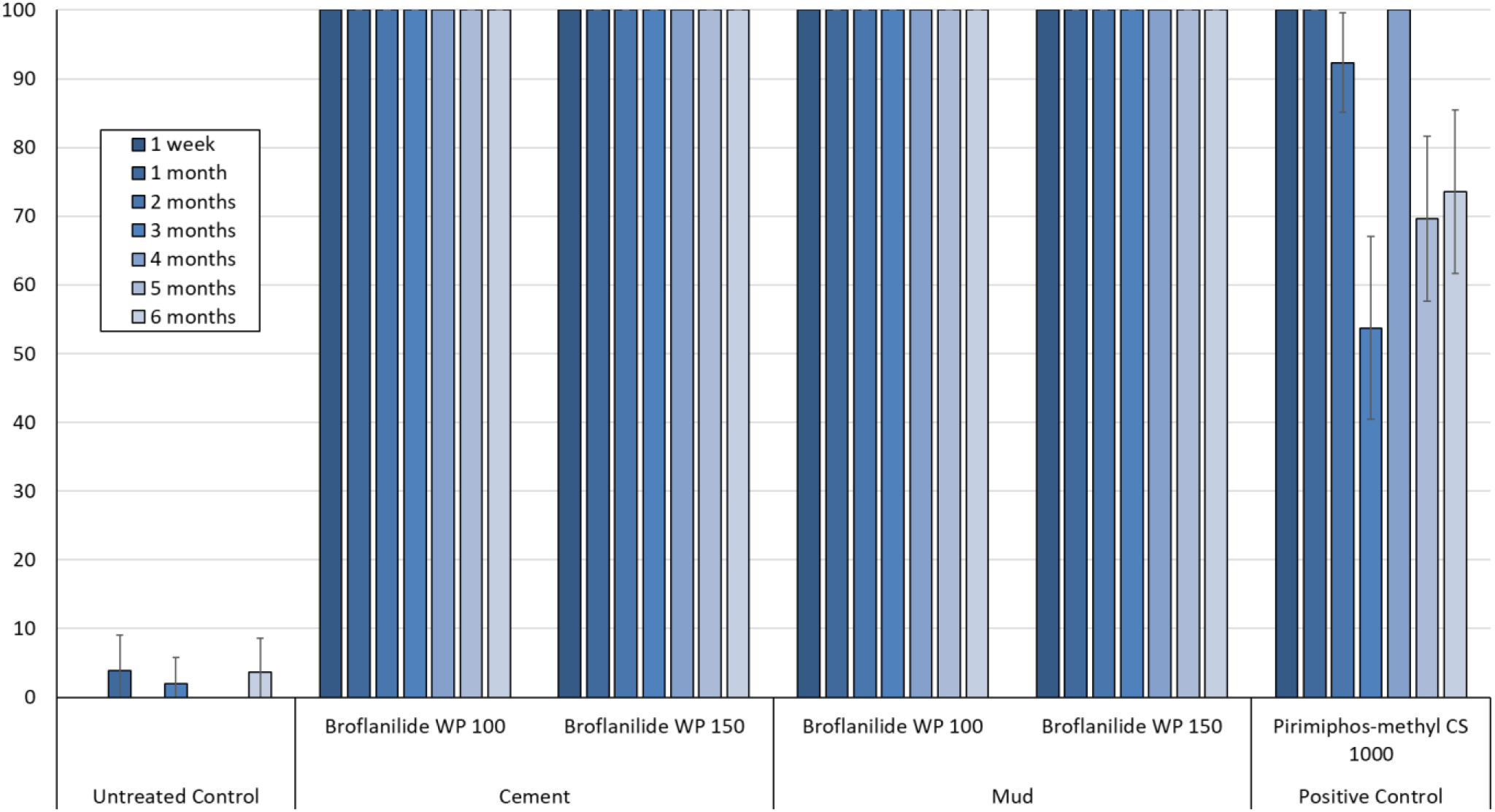
Cone bioassays mortality (72h) with pyrethroid-resistant *A. gambiae* s.l. Covè on Broflanilide WP-treated experimental hut walls. At each monthly interval, ∼50 2-5 days old female mosquitoes were exposed on treated of each hut in cohorts of 10 per cone. Error bars represent 95% confidence intervals.

Mortality rates achieved with broflanilide WP against *A. ziemanni* (Table 3) were generally higher than what was observed with *A. gambiae* s.l. (74%-88% vs. 50%-61%). Mortality in cement huts did not differ between the two application rates tested (74% with 100 mg/m^2^ vs 77% with 150 mg/m2 (P >0.05) but for mud-walled huts, mortality was significantly higher with the higher application rate of broflanilide WP (P<0.05). Mortality rates achieved with broflanilide WP on cement walls were also similar to what was observed with pirimiphos-methyl CS (P>0.05). Broflanilide WP also induced delayed mortality with *A. ziemanni* entering huts during the trial.

### Blood-feeding rates of wild pyrethroid-resistant A. gambiae s.l. in experimental huts

As expected of IRS treatments, blood-feeding rates of both mosquito species were generally very high across all huts (>90%). For pyrethroid-resistant *A. gambiae* s.l., there was no significant difference in blood-feeding rates between the two application rates (100 mg/m^2^ and 150 mg/m^2^) of broflanilide WP for either substrate type (P>0.05, Table 2). With *A. ziemanni*, while a significantly lower blood-feeding rate was observed with the higher dose of broflanilide WP in mud-walled huts, all other treatments tested gave similar blood-feeding rates irrespective of substrate type (Table 3).

### Mosquito mortality in cone bioassays on experimental hut walls treated with broflanilide WP is high and prolonged, lasting over 6 months

To assess the residual efficacy on the different hut wall substrates (mud and cement), monthly 30-min cone bioassays were performed using unfed pyrethroid-resistant *A. gambiae* s.l. Covè and insecticide-susceptible *A. gambiae* ss Kisumu strains. The results from the wall cone bioassays are presented in Figures 4 and 5. Broflanilide WP performed better on cement and mud wall substrates compared to Pirimiphos-methyl CS. For both strains, cone bioassay mortality with broflanilide WP remained over 80% with both doses and substrates throughout the 6-month trial while a drop below 80% was observed with pirimiphos-methyl CS within 2-4 months.

### Assessment of IRS application quality

To assess the quality of the IRS applications in the experimental huts, prior to spraying, filter papers measuring 5cm x 5cm were fixed on the hut walls to be sprayed as described in the WHO guidelines (*22*). After spraying, they were left to dry for 1 hour and then wrapped in aluminium foil and stored at 4^°^ C (+/-2°c) after which they were shipped within 2 weeks after IRS application to the Liverpool School of Tropical Medicine for chemical analysis by HPLC. The summary results showed that the IRS treatment applications rates were generally within an acceptable deviation of <50% from the target dose as recommended by WHO (*26*) (Table 4).

**Table 4.** Results for chemical analysis of filter papers from experimental huts treated with broflanilide WP and pirimiphos-methyl CS

| Treatment | Negative Control | Broflanilide WP |  |  |  | Pirimiphos-methyl CS |
| --- | --- | --- | --- | --- | --- | --- |
| Substrate | Cement | Cement |  | Mud |  | Cement |
| Target dose (mg/m<sup>2</sup>) | - | 100 | 150 | 100 | 150 | 1000 |
| Filter paper dosed (mg/m<sup>2</sup>) | 0 | 107 | 112 | 137 | 112 | 688 |
| 95% CI | - | [99 -115] | [103-122] | [129-145] | [95-130] | [95-130] |
| Deviation from target dose | - | +7% | -25% | +37% | -19% | -31% |

## Discussion

There is a critical need for new insecticides with novel modes of action for indoor residual spraying against malaria vectors. We investigated the bioefficacy of a newly discovered insecticide, broflanilide, (*17*) as an indoor residual treatment in WHO phase 1 laboratory bioassays and phase 2 experimental hut studies (*22*). Broflanilide binds to the γ-aminobutyric acid (GABA) receptor of the chloride channel at a different site to the cyclodiene insecticide dieldrin and the phenylpyrazole fipronil and thus presents a new mode of action for malaria vector control.

As a wettable powder formulation (VECTRON™ T500), broflanilide was tested for IRS against pyrethroid-susceptible and pyrethroid-resistant strains of *A. gambiae* s.l. on the principal wall substrates used in village housing in Benin. Both laboratory and experimental hut studies demonstrated the potency of this new mode of action when used for IRS. Mosquito mortality with broflanilide was high but slow compared to what is achievable with most neurotoxic public health insecticides, lasting up to 72 hours post-exposure. This delayed activity could be attributed to its mechanism of action; broflanilide has to be metabolized to its active form, desmethyl-broflanilide, before binding to its site of action (*18*). The small resistance ratio observed in CDC bottle bioassays (2.1) and high cone bioassay mortality rates (>80% for 6-14 months) achieved with the pyrethroid-resistant *An gambiae* s.l. strain from Covè - a strain which has shown >200 fold resistance to pyrethroids mediated by high kdr frequencies and overexpressed detoxifying P450 enzymes (*23*) - would indicate the absence of cross-resistance to broflanilide and pyrethroids. This demonstrates the potential of broflanilide WP to effectively control malaria vector populations that have developed intense resistance to pyrethroids.

In the experimental hut trial against wild pyrethroid-resistant vector mosquitoes, broflanilide applied at 100 mg/m^2^ and 150 mg/m^2^ induced 57-66% mosquito mortality for 6 months. As there was no difference in wild mosquito mortality between the two application rates tested, the lower dose of 100mg/m^2^ was chosen as a suitable IRS application rate for operational use of the insecticide in village communities.

Longer lasting IRS insecticide formulations are ideal for most endemic areas in Africa characterised by stable and extended malaria transmission as they offer continuous protection without the need for multiple resource-demanding and labour-intensive IRS campaigns. Earlier studies in Benin demonstrated the potential of the micro-encapsulated formulation of pirimiphos-methyl to provide improved and prolonged control of pyrethroid-resistant vector populations lasting 6-9 months when applied for on cement walls (*27*). This has been followed by several reports of significantly improved malaria control with one annual IRS campaign with pirimiphos-methyl CS in many epidemiological settings across Africa (*28-30*). In laboratory cone bioassays, mortality of susceptible and pyrethroid-resistant vector mosquito strains with broflanilide WP remained >80% for 6-14 months on mud and cement block substrates. The mortality achieved against wild free-flying pyrethroid-resistant mosquitoes in the experimental huts treated with broflanilide WP was >80% for >6 months demonstrating the potential of the insecticide to provide prolonged vector control and significantly improve malaria control in many endemic areas in Africa.

Although mutations in the GABA receptor conferring resistance to some non-competitive agonist agrochemicals such as cyclodienes and fipronil have been reported in malaria vectors across Africa (*31, 32*), the site of action of broflanilide within the GABA receptor has been demonstrated to be distinct from that of non-competitive antagonist (*20*). This suggests a low probability of cross-resistance to broflanilide in malaria vectors. Mutations in the GABA receptor which confer resistance to meta-diamides have however been detected in Drosophila (*20*). Studies to investigate the possible presence of this mutation and other mechanisms that could confer resistance to broflanilide in wild populations of malaria vectors will be advisable.

Despite its strong anthropophilic behaviour, a considerable number of *An gambiae* (81% *A. coluzzii* and 19% *An gambiae* ss), were attracted to the cow hosts during the experimental hut trial, although the numbers attracted were substantially lower than what would be expected in hut trials with human volunteer sleepers at the Covè experimental hut station (*14, 15*). The animal bait also attracted a more zoophilic Anopheline species into the experimental huts – *A. ziemanni*. Although this species has been much less studied compared to the *A. gambiae* complex, *A. ziemanni* has been previously implicated in malaria transmission in the North West region of Cameroon (*33*) and suspected to transmit malaria in Western Kenya (*34*). Such secondary vectors have been recognized for their importance in malaria transmission, as they may help to augment or extend the malaria transmission period. In our study, broflanilide killed 74%-88% of wild *A. ziemanni* entering the experimental huts over the 6-month trial thus demonstrating the potential of the insecticide to control this Anopheline species which could be sustaining malaria transmission as a secondary vector in some parts of Africa.

Among the strategies proposed by the GPIRM for mitigating the impact of insecticide resistance in malaria vectors, the rotation of IRS formulations containing insecticides with different modes of action is currently considered the most promising strategy for insecticide resistance management (*12*). The uptake of this strategy has been seriously limited by the very restricted number of safe and long-lasting IRS insecticides available to malaria control programmes (*35*). Two new IRS formulations containing clothianidin (a neonicotinoid) have recently been added to the WHO’s list of pre-qualified vector control products, and are already being deployed for IRS in many endemic countries (*13*). It is, however, crucial that vector control programmes do not become overly dependent on any one mode of action for IRS as this may lead to resistance evolving more rapidly. For the rotational strategy to work optimally, several modes of action need to be available at the same time to allow sub-national rotations and restrict selection of resistance to any single class of insecticide. Based on its novel mode of action and efficacy for IRS against pyrethroid-resistant malaria vectors as demonstrated in the present study, broflanilide WP shows potential to effectively complement other IRS formulations of insecticides in an IRS rotation plan that can manage insecticide resistance and extend the effective lives of these promising new insecticides.

## Conclusion

In this study, we demonstrate for the first time the efficacy of a wettable powder formulation of broflanilide (VECTRON™ T500), a newly discovered insecticide with a novel mode of action, for IRS against malaria vectors. VECTRON™ T500 showed high activity against both pyrethroid-susceptible and resistant strains of *A. gambiae* s.l. which lasted 6 months or more on local cement and mud substrates in both laboratory bioassays and experimental hut studies. Indoor residual spraying with broflanilide WP shows potential to provide improved and prolonged control of pyrethroid-resistant malaria vector populations.

## Materials and methods

### WHO Phase I Laboratory bioassays

#### CDC bottle bioassays to investigate resistance to broflanilide

The CDC bottle bioassays were performed with unfed 2-5 days old insecticide-susceptible *A. gambiae* s.s. Kisumu and pyrethroid-resistant *A. gambiae* s.l. adult female mosquitoes from Covè, Benin. Eight doses were selected between 5µg/bottle and 200µg/bottle as follows: 5µg, 10µg, 16.5µg, 27.1µg, 44.7µg, 73.7µg, 121.4µg and 200µg based on results from preliminary studies. Stock solutions were prepared by serial dilutions of the technical grade insecticide in acetone. 1 ml of each stock solution was used to coat each 250ml Wheaton bottle and 6 bottles were prepared per dose as described in the CDC bottle bioassay guideline (*36*). Approximately 150 female mosquitoes per insecticide dose were exposed for 1 hour in cohorts of 25 mosquitoes per bottle. Mosquitoes were also exposed to untreated control and alpha-cypermethrin 12.5µg treated bottles. Mosquitoes were held at 27 °C ± 2 °C and 80 ± 10% RH and mortality recorded after 72h based on preliminary evidence of a slower mode of action of broflanilide. Estimates of the dose required to kill 50% (LD50) and 95% (LD95) of each strain and the resistance ratio of the wild Covè strain relative to the susceptible Kisumu strain were generated by log dosage-probit analysis (PoloPlus version 1.0).

#### Preparation and treatment of block substrates

Cement and mud block substrates used in cone bioassays were moulded in Petri dishes (9cm diameter and 5mm thick) and dried at 27 °C ±2 °C and 80 ±10% RH for 30 days before insecticide application. Cement blocks were made by mixing cement with sand at a 1:1 ratio while mud blocks were made from local mud paste to which 10% cement was added to improve its durability and reduce cracking, in line with local practices. These substrates were treated using a Potter tower sprayer (Burkard Manufacturing Co Ltd) to achieve a homogeneous and accurate deposit of the target concentration of active ingredient (a.i) per unit area. Blocks were weighed before and after treatments to ensure the target amount of insecticide was delivered. All treated blocks were stored, unsealed at 30°C ± 2°C, 80% ± 10% RH in between bioassays. Four replicate blocks of each substrate-type were prepared for each dose of insecticide tested.

#### Dose-response cone bioassays

The dose-response cone bioassays were performed on mud and cement blocks treated with broflanilide WP at application rates of 5 mg/m^2^, 12.5 mg/m^2^, 25 mg/m^2^, 50 mg/m^2^ and 100 mg/m^2^, 1-week post block treatment using the insecticide-susceptible *A. gambiae* s.s. Kisumu strain. For each dose and substrate-type, forty (40) unfed 2-5 days old mosquitoes were exposed for 30 minutes in cohorts of 10 mosquitoes per con and per block. Mosquitoes were held at 27°C ± 2°C and 80% ± 10% RH post-exposure and knockdown was recorded after 1 hour. Mortality was recorded every 24 hours for up to 120 hours to investigate delayed mosquito mortality with broflanilide WP

#### Residual efficacy of broflanilide WP (VECTRON™ T500) in laboratory cone bioassays

For each insecticide dose and substrate-type used for assessment of residual efficacy, forty (40) unfed 2-5 days old female mosquitoes were exposed in cone bioassays for 30 minutes in cohorts of 10 mosquitoes per block. Mosquitoes were held under the same conditions as described earlier and mortality recorded every 24 hours for up to 72 hours. Bioassays were performed monthly for up to 6 months post-treatment and subsequently every two months for up to 18 months post-treatment.

### WHO Phase II experimental hut trial

#### Experimental hut site

The experimental hut site is located in an irrigated valley producing rice almost year-round and provides suitable breeding habitats for mosquitoes. The rainy season extends from March to October and the dry season from November to February. The vector population is resistant to pyrethroids and consists of both *An. coluzzii* and *A. gambiae* sensu stricto (s.s.) with the latter occurring at lower frequencies (∼23%) and mostly in the dry season. The local vector population in Covè is resistant to pyrethroids and DDT. The experimental huts used were of the West African design and are made from cement bricks with a corrugated iron roof. Each hut was built on a cement plinth surrounded by a water-filled moat to prevent the entry of scavenging ants and had a wooden framed veranda trap to capture exiting mosquitoes [23]. Mosquito entry occurred via four window slits each measuring 1 cm and situated on three sides of the hut.

#### Application of IRS treatments

The following treatments were tested in 6 experimental huts:

1. Untreated hut (negative control) – cement-walled hut
2. Broflanilide WP (VECTRON™ T500) applied at 100 mg/m^2^ – cement-walled hut
3. Broflanilide WP (VECTRON™ T500) applied at 100 mg/m^2^ – mud-walled hut
4. Broflanilide WP (VECTRON™ T500) applied at 150 mg/m^2^ – cement-walled hut
5. Broflanilide WP (VECTRON™ T500) applied at 150 mg/m^2^ – mud-walled hut
6. Pirimiphos-methyl CS (Actellic® 300CS) applied at 1000 mg/m^2^ – cement-walled hut

The IRS treatments were randomly allocated to experimental huts and were applied using a Hudson Xpert® compression sprayer equipped with a 8002 flat-fan nozzle and calibrated. To improve spray accuracy, spray swaths were marked before spraying and sprayed from the top to the bottom using the predetermined lance speed. After spraying each hut, the volume remaining in the spray tank was measured to assess the overall volume sprayed. All spray volumes were within 30% of the target.

#### Hut trial procedure

Six (6) cows were brought to sleep in the huts from 21:00 to 06:00 each trial night and were rotated between huts on successive nights to adjust for variation in individual attractiveness to mosquitoes. The cows were maintained according to institutional and national guidelines for the protection of experimental animals. The trial ran for 6 months from September 2018 to March 2019. Data collection was performed for 6 nights each week. On the 7th day of each week, the huts were cleaned and aired in preparation for the next cycle. Each morning, mosquitoes were collected from the room and veranda and brought to the laboratory where they were identified using standard identification keys and scored as fed or unfed and dead or alive. Live mosquitoes were provided with 10% glucose solution and mortality scored every 24 hours for up to 72 hours.

#### Outcome measures in experimental huts

The efficacy of each experimental hut treatment was expressed in terms of the following outcome measures:

- Exiting rates – the proportion of mosquitoes collected in the veranda
- Blood-feeding rates – the proportion of blood-fed mosquitoes
- Mortality – the proportion of mosquitoes found dead after a 120-hour holding time

#### Monthly residual cone bioassays on treated experimental hut walls

Five cones were fixed to each treated surface of each hut (1 per wall + 1 on the ceiling). Approximately fifty (50) unfed 2-5 days old mosquitoes of each strain were exposed for 30 minutes to each hut treatment in batches of 10 mosquitoes per cone. After exposure, mosquitoes were transferred into netted plastic cups and supplied with 10% sugar solution. Mortality was recorded every 24 hours up to 72 hours.

### Data management and statistical analysis

Cone bioassay data were pooled for each substrate and dose and mean mortalities obtained using Microsoft Excel. CDC bottle bioassay data were analysed using log dosage-probit mortality analysis (Poloplus version 1.0) to determine the lethal concentration (LC) required to kill 50% (LC50) and 95% (LC95) of exposed mosquitoes. The diagnostic dose was defined as twice the LC95 for the insecticide-susceptible Kisumu strain while the resistance ratio of the wild field-collected pyrethroid-resistant *An gambiae* s.l. Covè strain was obtained by dividing its LC50 by that of the susceptible Kisumu strain. Proportional outcomes (blood-feeding, exophily and mortality) for each experimental hut treatment were analysed using blocked logistic regression in Stata version 15.1 with adjustments for the attractiveness of the individual cow hosts.

### Ethical considerations

Institutional ethical approval for the study was obtained from the Ethics Review Committee of the Ministry of Health Benin (Ethical decision n°39). The cows used in experimental huts to attract mosquitoes were maintained following institutional standard operating procedures (SOPs) designed to improve care and protect animals used for experimentation. During the day, the cows were allowed to graze freely in an open field not too far from the experimental hut site. A veterinarian was available throughout the study to examine the cows daily and any cows found unwell was replaced and treated appropriately.

## Declarations

### Consent for publication

Not applicable

#### Availability of data and material

The datasets used and/or analysed during the current study are available from the corresponding author on reasonable request.

## Authors’ contributions

CN co-designed the study, supervised the project, analysed the data and prepared the final manuscript. RG supervised the hut trial and contributed to data analysis and manuscript preparation. EV performed the laboratory bioassays while AF performed the hut trial. TS contributed to data analysis. MA provided administrative and logistics support. MR co-designed the study and contributed to manuscript preparation.

## Acknowledgements

We thank Dr. Kunizo Mori of Mitsui Chemicals Agro, Inc for supplying the insecticide. We also thank the technical staff of CREC (Abibath Odjo, Damien Todjinou, Josias Fagbohoun, Martial Gbegbo etc) for their assistance. We appreciate Dr. Graham Small, Dr. Derric Nimmo and Dr. Sarah Rees of IVCC for coordination. We acknowledge Dr. Mark Paine of LSTM for performing the chemical analysis. We are grateful to the rice farmers at Cove for their support in the hut study.

## Authors’ information

Not applicable

## List of abbreviations

IRS: Indoor Residual Spraying
WHO: World Health Organization
PQ: Prequalification team
GPIRM: Global Plan for Insecticide Resistance Management
WHOPES: WHO Pesticide Evaluation Scheme
CREC: Centre de Recherche Entomologique de Cotonou
LSHTM: London School of Hygiene & Tropical Medicine
Kdr: Knockdown resistance

